# Methamphetamine Self-Administration Differential Effects on Mesolimbic Glutathione Levels, Mitochondrial Respiration and Dopamine Neuron Firing Activity

**DOI:** 10.1101/2022.09.12.507550

**Authors:** Sergio Dominguez-Lopez, Bumsoo Ahn, Kavithalakshmi Sataranatarajan, Rojina Ranjit, Pavithra Premkumar, Holly Van Remmen, Michael J. Beckstead

## Abstract

Acute and neurotoxic regimens of METH are known to increase reactive oxygen species (ROS), affect redox homeostasis, and lead to cellular damage in dopamine neurons. However, functional changes induced by long-term METH self-administration on mitochondrial respiratory metabolism and redox homeostasis are less known. To fill this gap in our knowledge, we implanted adult mice with a jugular catheter and trained them to nose poke for METH infusions in operant chambers. After completing several weeks of METH self-administration exposure, we collected samples of the ventral striatum (vSTR) and the ventral midbrain (vMB), containing the nucleus accumbens (NAc) and the ventral tegmental area (VTA), respectively. We used HPLC to determine the levels of the ROS scavenger glutathione in its reduced (GSH) and oxidized (GSSG) forms. Then, we used high-resolution respirometry to determine the oxygen consumption rate (OCR) of mitochondrial complexes under several substrates and inhibitors. Finally, we used in vivo single-unit extracellular recordings to assess changes in dopamine neuron firing activity in the VTA. METH self-administration produces a progressive decrease of the GSH pool in vST, which correlates with METH lifetime intake. We observed increased mitochondrial respiration across the two mesolimbic regions, but only vMB OCR correlates with METH lifetime intake. We recorded an increased number of spontaneously active dopamine neurons with decreased firing rate and burst activity in the VTA. METH lifetime intake inversely correlates with firing rate, the percentage of spikes in a burst, and directly correlates with the number of neurons per track. We conclude that METH self-administration progressively decreased the antioxidant pool in sites of higher dopamine release and produced an increased mitochondrial metabolism in the mesolimbic areas, probably derived from the increased number of dopamine neurons actively firing. However, dopamine neuron firing activity is decreased by METH self-administration, reflecting a new basal level of dopamine neurotransmission in response to the prolonged effects of METH on dopamine release and circuitry feedback.

## Introduction

The functional and metabolic changes in the brain of drug users transitioning from recreational to constant drug use and addiction are not entirely understood. However, it is well established that the dopamine mesolimbic pathway directly or indirectly mediates the rewarding, reinforcing, and addictive properties of methamphetamine (METH) and other used substances (Nestler and Luscher, 2019). Indeed, used drugs by humans can increase not only extracellular dopamine release in axon terminal areas such as the nucleus accumbens (NAc) and the prefrontal cortex but also in somatic areas such as the ventral tegmental area (VTA) (Bhimani et al., 2021; Di Chiara and Imperato, 1988; Kalivas and Duffy, 1988).

The acute reinforcing effects of METH in the brain derive from this ability to increase extracellular dopamine. Due to its monoaminergic structure, the plasma membrane protein responsible for the clearance and reuptake of dopamine after its release (the dopamine transporter, DAT) can recognize and use METH as a substrate (Han and Gu, 2006; Kuczenski et al., 1995). METH and other amphetamines can reverse-transport dopamine to the extracellular space, a process known as efflux (Fleckenstein et al., 2007). Inside the cell, METH affects monoamine homeostasis causing an increase in the activity of the enzyme synthesizing dopamine (tyrosine hydroxylase, TH) and inhibiting the action of the enzyme that metabolizes it (monoamine oxidase, MAO) (Hotchkiss and Gibb, 1980; Suzuki et al., 1980). METH also disrupts the function of the vesicular monoamine transporter (VMAT2), inhibiting vesicular storage and promoting an increase in cytoplasmic dopamine (Brown et al., 2000; Rau et al., 2006). However, the effects of METH on these enzymes are region specific and depend on the dose, frequency, and exposure time (Fibiger and Mogeer, 1971; Haughey et al., 1999; Moreira da Silva Santos et al., 2019). Following injection, METH plasma concentrations peak within a few minutes and have a long half-life (about 10 hours) which contributes to its addictive properties and produces neurotoxic effects (Cruickshank and Dyer, 2009).

Besides modifying dopamine synthesis and release, we are beginning to understand metabolic changes in mesolimbic circuitry after long-term METH use. One property of METH exposure is the generation of oxidative stress primarily derived from increased dopamine catabolism by mitochondria-bound MAO (Graves et al., 2020; Yang et al., 2018). The supraphysiological degradation of dopamine could then unbalance redox metabolism and mitochondrial function by increasing the production of reactive oxygen species (ROS), hydrogen peroxide (H_2_O_2_), and superoxide anions (O_2-_) (Quinton and Yamamoto, 2006; Yang et al., 2018). Changes in the ratio of reduced (GSH) to oxidized (glutathione disulfide, GSSG) glutathione is a well-established marker of oxidative stress, and GSH functions as a ROS scavenger (Aoyama et al., 2008; Uys et al., 2014). Interestingly, decreased glutathione levels in the caudate have been observed postmortem in METH users with higher dopamine content loss (Mirecki et al., 2004). In rats, high-dose binge regimens of METH can deplete striatal glutathione (Moszczynska et al., 1998), and there is evidence that the glutathione redox system could be of clinical relevance to protecting against METH-induced oxidative damage (Barayuga et al., 2013). However, the effects of ongoing METH use on mesolimbic glutathione levels are not currently known.

METH exposure also leads to functional alterations of the mitochondrial electron transport chain and decreases the expression of some mitochondrial complex subunits (Bazylianska et al., 2021; Brown et al., 2005). This adds to the capability of excessive dopamine oxidation to alter mitochondrial respiration (Berman and Hastings, 1999). However, most research on METH, dopamine, and mitochondrial function has been done in the context of neurotoxic degeneration of the substantia nigra (SN). Only recently has it become a point of interest for substance use research (Bazylianska et al., 2021; Graves et al., 2021). To our knowledge, there are no studies related to how METH self-administration affects mitochondrial respiratory function. However, some studies showed that noncontingent METH induces an acute increase in ATP consumption and disrupts mitochondrial respiratory function in dopamine mesolimbic areas (Burrows et al., 2000; Shiba et al., 2011). This finding is important because dopamine neurons are vulnerable to mitochondrial dysfunction due to their high energy demands, unmyelinated axonal arborization, and continuous firing activity (Ni and Ernst, 2022; Pacelli et al., 2015; Pissadaki and Bolam, 2013). Indeed, the relationship between mitochondria metabolism and neuronal firing is essential for the maintenance of brain function (Ruggiero et al., 2021). For example, deletion of the mitochondrial transcription factor A in dopamine neurons increases the variability of firing in the MitoPark mouse model of nigral degeneration (Branch et al., 2016; Good et al., 2011) and disruption of mitochondrial complex I in SN dopamine neurons slowed their autonomous pacemaker firing pattern and trigger parkinsonian-like degeneration (Gonzalez-Rodriguez et al., 2021). Therefore, one may predict alterations in dopamine neuron firing to develop in parallel to mitochondrial dysfunction following METH exposure, but this remains understudied.

This study aimed to determine the sensitivity of dopamine-rich mesolimbic regions to unbalanced mitochondrial redox homeostasis induced by prolonged METH self-administration. First, we trained and exposed mice to long-term METH self-administration and collected samples of the ventral striatum (vST) and the ventral midbrain (vMB) to determine changes in the balance of GSH and GSSG. We then assessed the function of mitochondrial complexes by determining their oxygen consumption rates (OCR) and H_2_O_2_ production. Finally, we performed *in vivo* electrophysiology to determine changes in firing activity (basal tonic and burst firing activity) of VTA dopamine neurons after long-term exposure to METH-self administration. Our study contributes to the small but growing literature on mitochondria in substance use disorder and motivation.

## Materials and Methods

### Animals

We used adult (five to seven-month-old) C57BL/6J male mice (Jackson labs, Bar Harbor, ME), which were group-housed in polycarbonate boxes with rodent bedding and shredding material and kept on a 12/12-hour reverse light cycle (lights off at 0900 h) with *ad libitum* access to food and water. The Institutional Animal Care and Use Committee at the Oklahoma Medical Research Foundation revised and approved all procedures in this study.

### Drugs

We dissolved METH hydrochloride (generously provided by the National Institute on Drug Abuse drug supply program) and chloral hydrate (Sigma-Aldrich) in sterile saline (0.9 % NaCl).

### Catheter implantation

We implanted mice with an indwelling catheter in the right jugular vein as previously described (Dominguez-Lopez et al., 2018). After surgery, we housed the mice individually and allowed them at least seven days to recover. We flushed the catheters daily with 0.02 ml heparinized saline (30 IU/ml) beginning three to four days after placement. We assessed catheter patency by flushing with sterile saline before each operant session and at the end with heparinized saline.

### Operant self-administration

We conducted two-hour operant sessions as previously described (Dominguez-Lopez et al., 2018). The modular mouse operant chambers had two nose poke holes (Lafayette Instruments, Lafayette, IN), one illuminated from the inside by a dim green light. During training, we rewarded responses in the illuminated hole on a fixed ratio of 1 (FR1) for infusions 1 to 5, FR2 for infusions 6 to 7, and FR3 for the remainder of the session. Upon completing the response requirement, the green light stimulus turned off, initiating a 15-second timeout, and METH was delivered intravenously (0.1 mg/kg/infusion, 12 µl) over 2 seconds, accompanied by a 2 kHz tone. METH infusion dosage was calculated based on the average weight of the mice (32 gr) and corrected by individual body weight to obtain intake (mg/kg/session). Responding during the timeout was recorded but not reinforced. We considered self-administration learned when the number of infusions earned was equal or higher than eight and the number of responses in the correct hole was ≥ 70% of the total responses for at least two consecutive days.

### Stabilization, extinction, and progressive ratio procedures

Following training, we kept the mice in an FR3 schedule to stabilize their METH intake. We then placed the mice on an extinction schedule for ten days (EXT), during which we attached the mice to the intravenous tether without delivering infusions or cues (light or sound) during the 2-hour session. EXT was followed by one day on an FR3 cue-induced reinstatement (Cue-R) schedule (nose pokes resulted in the sound cue, but not METH infusion). Then, we returned the mice to the FR3 schedule to finally test them with a five-hour session on a progressive ratio schedule (PR). During PR sessions, the number of nose pokes necessary for an infusion was gradually increased (1, 2, 4, 6, 9, …) following the procedure described by Richardson and Roberts (1996).

### Tissue collection for OCR and glutathione determinations

We collected our fresh tissue samples by adapting a technique used for *ex vivo* electrophysiology (Howell et al., 2020). Briefly, we anesthetized mice with isoflurane before dissection of their brain and immediately placed it in an ice-cold oxygenated cutting solution containing the following (in mM): 250 sucrose, 26 NaHCO_3_, 2 KCl, 1.2 NaH_2_PO_4_, 11 glucose, 7 MgCl_2_, 0.5 CaCl_2_. We collected coronal brain slices (200 μm) using a VT1200S vibratome (Leica, Deer Park, IL). We carefully dissected the sections containing the vST and the vMB with a stereoscope, limiting tissue sampling to the NAc and VTA. We collected and processed one or two samples per region per animal. After dissection, we kept the samples in the ice-cold cutting solution before transferring to Buffer X medium for OCR / H_2_O_2_ production rate measurements on the same day or directly frozen them in liquid nitrogen and stored them at -80 ºC for glutathione determinations.

### HPLC separation and electrochemical detection (ECD) of glutathione

We separated GSH and GSSG using a Dionex UltiMate 3000 HPLC-ECD 3000RS system (Thermo Scientific, Waltham, MA) with an Accucore RP-MS (150 mm X 2.1 mm) reverse-phase column (Thermo Scientific, Waltham, MA). We used previously reported HPLC conditions (Ahn et al., 2019) with slight modifications, including a mobile phase of 25 mM sodium phosphate, 0.5 mM 1-octane sulfonic acid, and 4 % acetonitrile (adjusted to pH 2.7 using *o*-phosphoric acid) with 35 ºC column temperature and 0.5 mL/min flow rate. First, we used 5 % of meta-phosphoric acid to homogenize pulverized frozen brain tissue samples. Then, we centrifuged the samples at 16400 rpm for 15 minutes at 4 ºC and transferred the supernatant to HPLC autosampler vials. One aliquot of the supernatant was used for protein estimation by the BCA method according to the manufacturer’s instructions (Pierce, Rockford, IL, USA). Before we injected the samples, the system was equilibrated to maintain a stable baseline. After every run, we apply a clean cell potential of 1900 mV for 10 s. We used a conditioning coulometric OMNI cell (6020RS, Thermo Scientific) before the Boron-doped diamond electrode to chromatographically filter electrochemically active compounds that could interfere with GSH and GSSG detection. The GSH and GSSG peak eluted at 1.14 min and 1.98 min using 500 mV and 1200 mV for OMNI and BDD, respectively. The concentration peaks were quantified using Chromeleon CDS Chromatography software (Thermo Scientific).

### Simultaneous assessment of OCR and H_2_O_2_ generation rate

We followed a previously published protocol with some modifications for brain tissue (Ahn et al., 2019). Brain samples were transferred to ice-cold Buffer X medium, containing (in mM) 7.23 K_2_EGTA, 2.77 CaK_2_EGTA, 20 imidazole, 0.5 DTT, 20 taurine, 5.7 ATP, 14.3 PCr, 6.56 MgCl_2_-6H_2_O, 50 K-MES (pH 7.1). We weighed one or two brain samples (∼2-5 mg). We permeabilized them in Buffer X with saponin (50 µg/mL) for 30 mins, followed by 3 × 5 minutes washes in ice-cold Buffer Z containing (in mM): 105 K-MES, 30 KCl, 10 K_2_HPO_4_, 5 MgCl_2_-6H_2_O, 0.5 mg/ml Bovine Serum Albumin (BSA), 0.1 EGTA (pH 7.1). OCR and the rate of mitochondrial H_2_O_2_ production were simultaneously determined using high-resolution respirometry with an Oxygraph-2k (O2k, Oroboros Instruments, Innsbruck, Austria). OCR was determined using an oxygen probe, while rates of H_2_O_2_ generation were determined using the O2k-Fluo LED2-Module Fluorescence-Sensor Green. Measurements were performed on permeabilized fibers in Buffer Z media at 37 ∼C containing: 10 µM Amplex UltraRed (Molecular Probes, Eugene, OR), 1 U/mL horseradish peroxidase (HRP), and 25 µM blebbistatin. HRP catalyzes the reaction between hydrogen peroxide and Amplex UltraRed to produce the fluorescent resorufin (excitation: 565 nm; emission: 600 nM). Each day of experiments, we established a standard curve to convert the fluorescent signal to nanomolar H_2_O_2_. We subtracted the background resorufin production from each measurement. OCR and H_2_O_2_ production were determined using sequential additions of substrates and inhibitors as follows: glutamate (GLU,10 mM), malate (MAL, 2 mM), Adenosine diphosphate (ADP, 5 mM), succinate (SUC, 10 mM), rotenone (ROT, 1 µM), Antimycin A (AA, 1 µM), and N, N, N’, N’-tetramethyl-p-phenylenediamine (TMPD, 0.5 mM) immediately followed by ascorbate (ASC, 5 mM). All respiration measurements were normalized to AA to account for non-mitochondrial residual oxygen consumption. We calculated the oxidative phosphorylation (OXPHOS) capacity for complexes I & II using the formula 1-C_p_/C_l_, where C_p_ corresponds to complex-linked active phosphorylation respiration (after ADP injection), and C_l_ equals leak state respiration (GLU/MAL). We normalize data for both OCR and rates of H_2_O_2_ generation by milligrams of tissue wet weights and analyze with the DatLab software (Oroboros Instruments, Innsbruck, Austria).

### Single-unit In Vivo extracellular recordings

We anesthetized mice with chloral hydrate (400 mg/kg, i.p.) and placed them in a stereotaxic apparatus (Kopf Instruments, Tujunga, CA), drilling a hole in the skull to give access to the brain, as previously described (Dominguez-Lopez et al., 2021). Using an infrared heating lamp, we maintained the mice’s body temperature at 35-36.5 °C. Single barrel glass micropipettes with an internal diameter of 1.5 mm (Sutter Instruments, Novato, CA) were pulled to a shank length of approximately 0.6 cm (5-8 MΩ) using a Narishige PC-10 puller (Tokyo, Japan) and filled with 2% Pontamine sky blue dye (Alfa Aesar, Haverhill, MA) dissolved in sodium acetate (0.5 M, pH 7.5). We lowered the electrodes into the VTA using a hydraulic microdrive (Stoelting, Wood Dale, IL) to the following stereotaxic coordinates: -2.9 to -3.6 mm from Bregma, ± 0.2 to ± 0.7 mm from the midline, and -3.5 to -5.5 mm from the brain surface (Franklin and Paxinos 2008). We performed one to five electrode descents per mouse, recording the spontaneous electrical activity of single cells using an MDA-4 amplifier system (Bak Electronics, Umatilla, FL). Spike frequencies and waveforms were collected and stored on a computer using Spike 2 and a Micro1401-3 acquisition unit (CED, Cambridge, UK). We identified dopamine neurons based on well-established electrophysiological properties: a broad action potential (> 2.5 ms), biphasic or triphasic waveform, and slow firing (0.2-12 Hz) (Dominguez-Lopez et al., 2014; Ungless and Grace, 2012). We measured neuronal activity by calculating the mean firing frequency, expressed as the number of spikes per second (Hz). Also, we analyzed the number of spontaneously active neurons per electrode descent or track as an index of population activity (Ungless and Grace, 2012).

Additionally, the burst activity of dopamine neurons was analyzed using a Spike 2 script based on published criteria (Dominguez-Lopez et al., 2014; Ungless and Grace, 2012). We defined a burst as a train of at least two spikes with an initial interspike interval (ISI) ≤ 80 ms and a maximum ISI of 160 ms within a regular low-frequency firing pattern and decreased amplitude from the first to the last spike within the burst. On the final track for each mouse, Pontamine sky blue dye was injected iontophoretically by passing a constant positive current of 20 μA for 5-10 min through the recording pipette to mark the recording site. Mice were then decapitated, and their brains extracted and placed in paraformaldehyde solution (4%) for at least two days. Finally, we localized the labeled site by visually inspecting 40 μm coronal sections.

## Statistical Analysis

We analyzed our data using SigmaStat 4.0 statistical suite (Systat, San Jose, CA). We used the Student t-test for comparisons between Naive and METH self-administration groups. Two-way ANOVA was used to determine interactions between brain regions and METH exposure status. We used Bonferroni-corrected t-tests for *post hoc* comparisons after ANOVAs. Finally, we used Pearson product-moment (r) to determine the relationships between METH intake with other variables. We report the data as the mean ± standard error or as the data distribution in box plots. Statistical values of p ≤ 0.05 were considered significant.

## Results

We implanted 35 mice with intravenous catheters and trained them to nose poke for METH infusions (Figure 1A). We excluded only three mice from our study since they developed catheter patency issues and never fulfilled our criteria for learning self-administration. Figure 1B shows the self-administration learning curve and stabilization of the FR3 schedule. Our mice were exposed to METH in an average of 37 operant sessions (range: 24 to 53 sessions, excluding EXT and Cue-R sessions) over 66 calendar days (range: 33 to 100 days). METH self-administration dosage across FR3 sessions was 1.66 ± 0.05 mg/kg (Figure 1C) and lifetime cumulative intake was 61.88 ± 8 mg/kg (range: 15.67 to 188.6 mg/kg). METH intake during FR3 sessions in our mice was above the estimated recreational dose of METH (0.4 mg/kg) and fell within the estimated intake of chronic users in a day (1.4 - 14 mg/kg) (Moszczynska and Callan, 2017). Our mice extinguished their behavior when METH infusions were unavailable (EXT, Figure 1B), displayed METH seeking in the presence of cues (Cue-R, Figure 1D), and showed motivation to work for METH infusions in the PR schedule (Figure 1D). METH intake directly correlated with the response observed in PR (r = 0.56, p =0.001, Figure 1E) tests.

**Figure 1.**
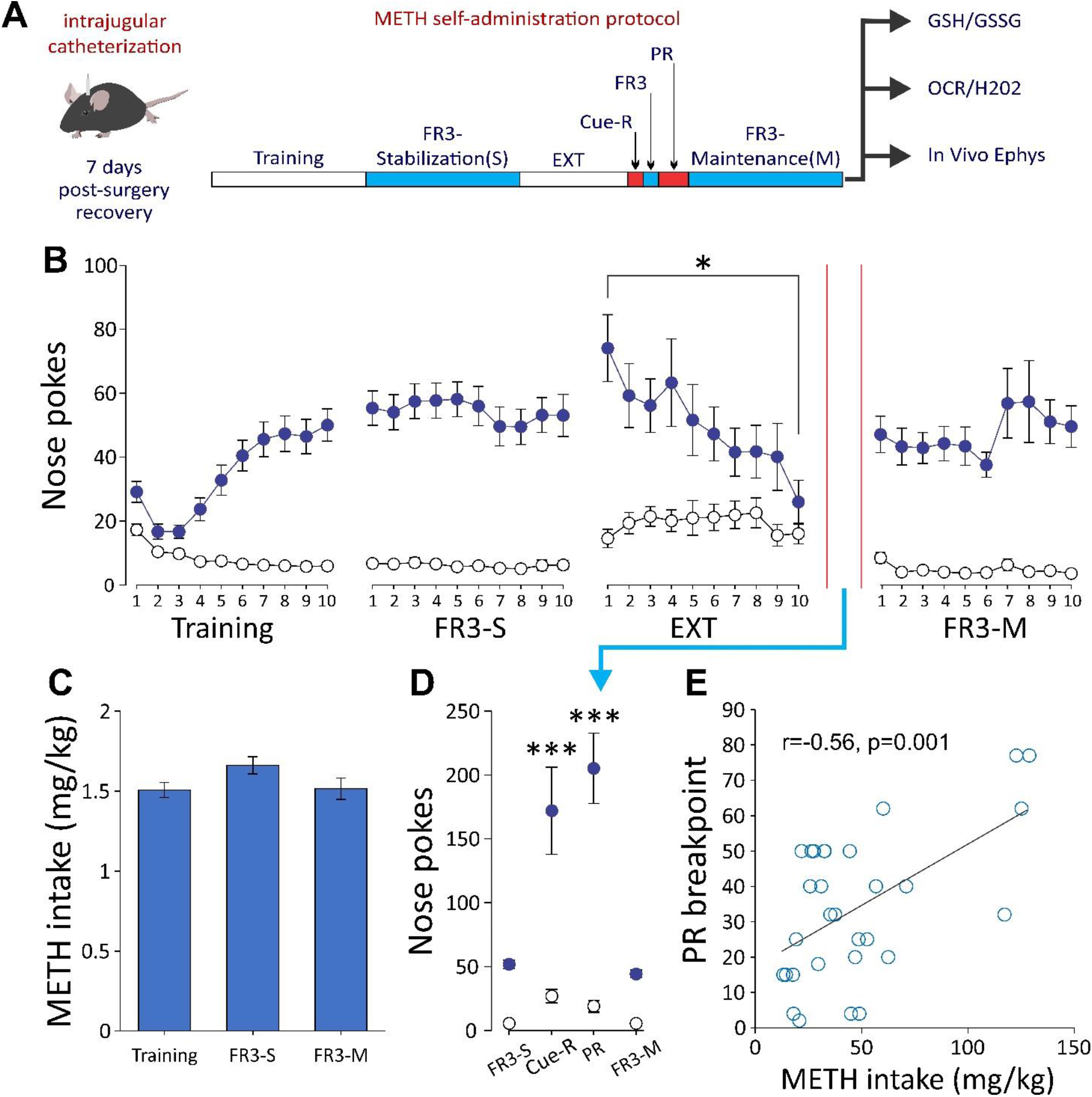
Methamphetamine (METH) self-administration in mice. A) General procedure in which mice were trained showing the different phases; training, fixed ratio 3 for behavior stabilization (FR3-S), extinction (EXT), Cue-reinstatement (Cue-R), Progressive Ratio (PR), and FR3 for maintenance until endpoint (FR3-M). The determinations that followed are indicated; HPLC reduced glutathione (GSH) and oxidase glutathione (GSSG), high-resolution respirometry for oxygen consumption rate (OCR) and peroxide (H_2_O_2_) production rate, and in vivo electrophysiology (Ephys) of single-dopamine neuron firing activity. B) Plots indicating the number of nose pokes in the hole that delivered METH infusion (blue) and in the inactive wrong hole (open circles) across the different tests. The red lines indicate when EXT, Cue-R, and PR took place). C) METH intake across the training and FR3 sessions was constant. D) Responses in the active nose poke hole on Cue-R and PR test days increased above the average of the last 3 FR3-S and the first three FR3-M sessions. E) The lifetime amount of METH intake before the PR test directly correlates with the maximum ratio completed or breakpoint. Bonferroni Student-t test after ANOVA: *** p <0.001 (vs. FR3-S and FR3-M); Simple t test: * p < 0.05.

### METH self-administration selectively decreases GSH levels in vST

First, to test the impact of METH self-administration on redox homeostasis along the mesolimbic pathway, we quantified the levels of GSH and GSSG using HPLC with ECD. We also calculated the GSH/GSSG ratio, an index of redox potential in several tissues and pathological brain conditions (Ahn et al., 2019; Gawryluk et al., 2011; Gu et al., 2015). We collected samples from nine mice exposed to METH self-administration (within two days after the last self-administration session) and seven naive mice.

We detected an interaction of METH self-administration and brain region for GSH levels (F(1,29) = 7.57, p = 0.01). In naive conditions, the vST levels of GSH were higher than in vMB (p = 0.009, Figure 2A), but vST GSH decreased in the METH self-administration group (p < 0.01). We detected only a trend towards lower levels of GSSG in METH exposed animals independently of brain region (F(1,29) = 3.6, p =0.067, Figure 2B) and a lower GSH/GSSG ratio in vMB samples independently of METH exposure (F(1,29) = 4.71, p = 0.038, Figure 2C). Our results suggest that the vST has a higher oxidative buffer capability than the vMB due to a higher basal level of GSH. Interestingly, the lifetime METH intake of the individual mice inversely correlates to both GSH (r = -0.561, p = 0.02) and GSSG (r = -0.512, p = 0.03) levels but not for the GSH/GSSG ratio (r = 0.348, p = 0.17) in vST samples (Figure 2D). This linear correlation was not observed in vMB samples (Figure 2D), suggesting that METH self-administration preferentially alters glutathione redox homeostasis at sites of high dopamine release, such as the NAc.

**Figure 2.**
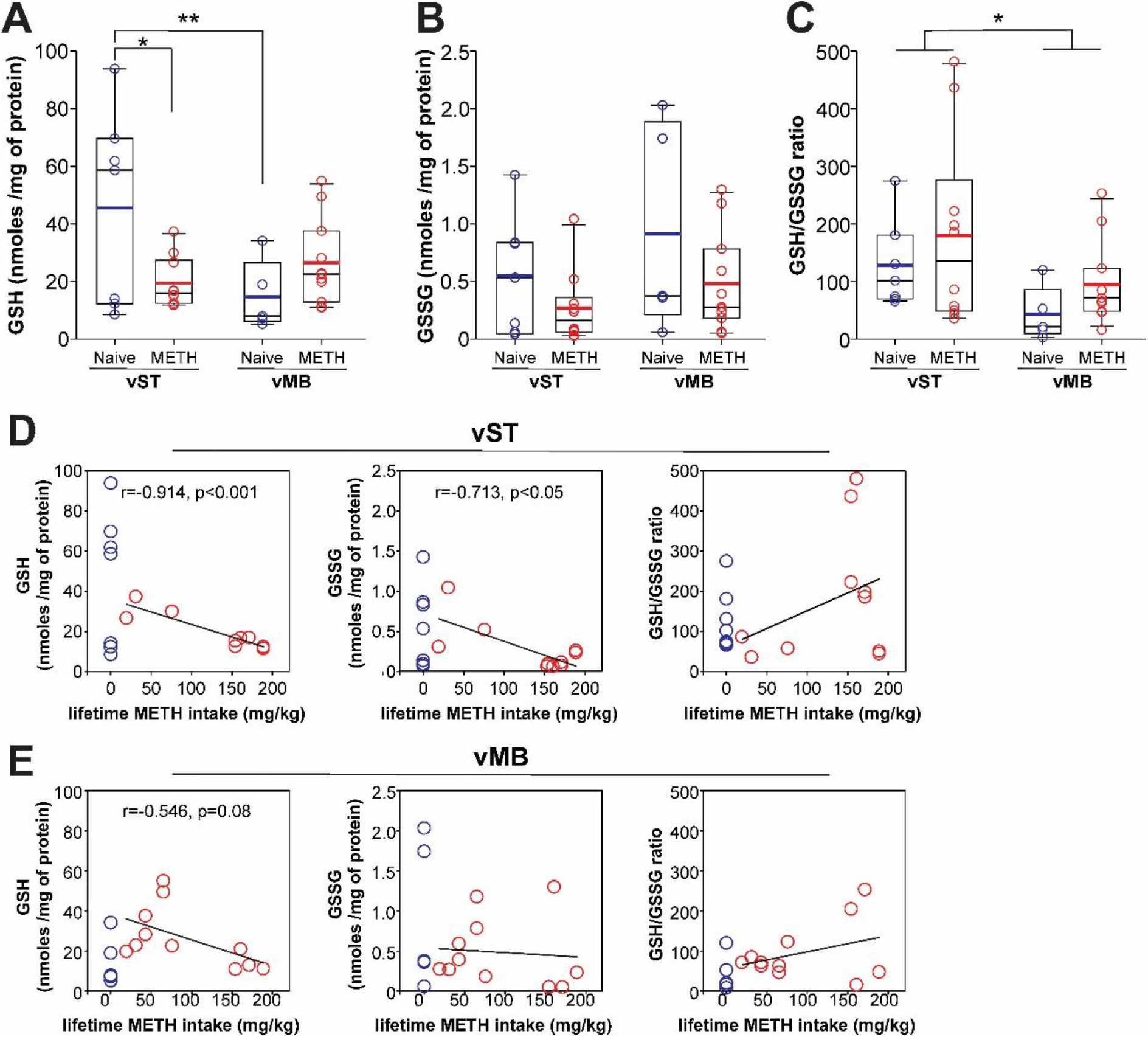
Levels of the antioxidant glutathione in the ventral striatum (vST) and the ventral midbrain (vMB) after methamphetamine (METH) self-administration. A) The levels of the reduced form of glutathione (GSH) are higher in the vST of drug-naive mice (Naive) compared with vMB levels. In mice exposed to METH self-administration, GSH levels are reduced in VST but not vMB. B) No significant changes were observed in the levels of the oxidized form of glutathione (GSSG) between regions or compared after METH self-administration. C) The antioxidant buffer capability of glutathione, assessed as the GSH/GSSG ratio, was lower in vMB compared with vST samples, independent of METH exposure status. D) Lifetime METH intake inversely correlated with levels of GSH and GSSG in the vST, E) but not in the vMB. No significant correlation was detected for the GSH/GSSG ratio. Bonferroni Student-t test after ANOVA: * p < 0.05, **p < 0.01.

### METH self-administration increases mitochondria OCR in mesolimbic regions

Next, we tested the impact of prolonged exposure to METH self-administration on mitochondria function. We used high-resolution respirometry to determine the OCR of mitochondrial complexes and their corresponding production of H_2_O_2_. We used permeabilized vST and vMB sections to preserve local circuit dynamics lost using dissociated cells or isolated mitochondria (Dias et al., 2018). We collected samples from 14 mice exposed to METH self-administration and 12 naive mice. Most tissue samples from METH-exposed mice were collected a day after the last self-administration session. Two mice were processed two days after the final session. Figure 3A shows an example of our basic protocol for high-resolution respirometry. Basal tissue OCR without reagents was higher in samples from METH-exposed mice independently of the brain region (F(1,50) = 6.103, p = 0.017, Figure 3B). In a similar way, after correcting for non-mitochondrial respiration (AA response), increased OCR was observed in samples from mice with METH self-administration exposure in all conditions: GLU/MAL (F(1,50) = 6.879, p = 0.012, ADP (F(1,50) = 5.869, p = 0.019), SUC (F(1,50) = 6.835, p = 0.012, ROT (F(1,50) = 12.084, p = 0.001) and ASC/TMPD (F(1,50) = 11.072, p = 0.002), as it is shown in Figure 3C. We interpreted these data as an overall increase in mitochondria respiration in dopamine mesolimbic regions due to METH self-administration exposure. Interestingly, respiration after AA application remained at a higher rate in vMB samples compared with vST samples (F(1,50) = 4.021, p = 0.05), with a trend of METH exposed tissue to also maintain higher OCRs after AA application (F(1,50) = 3.08, p = 0.08, Figure 3D). We can translate the difference in complex III response to inhibition by AA as increased non-mitochondrial respiration in the vMB. This difference may also reflect less complex III content or subunit composition in vMB versus vST neurites (Kilbride, Gluchowska et al. 2011, Bazylianska, Sharma et al. 2021). Further data analysis indicates that METH intake directly correlated to OCR in vMB in almost all conditions but not with OCR from vST samples (data not shown).

**Figure 3.**
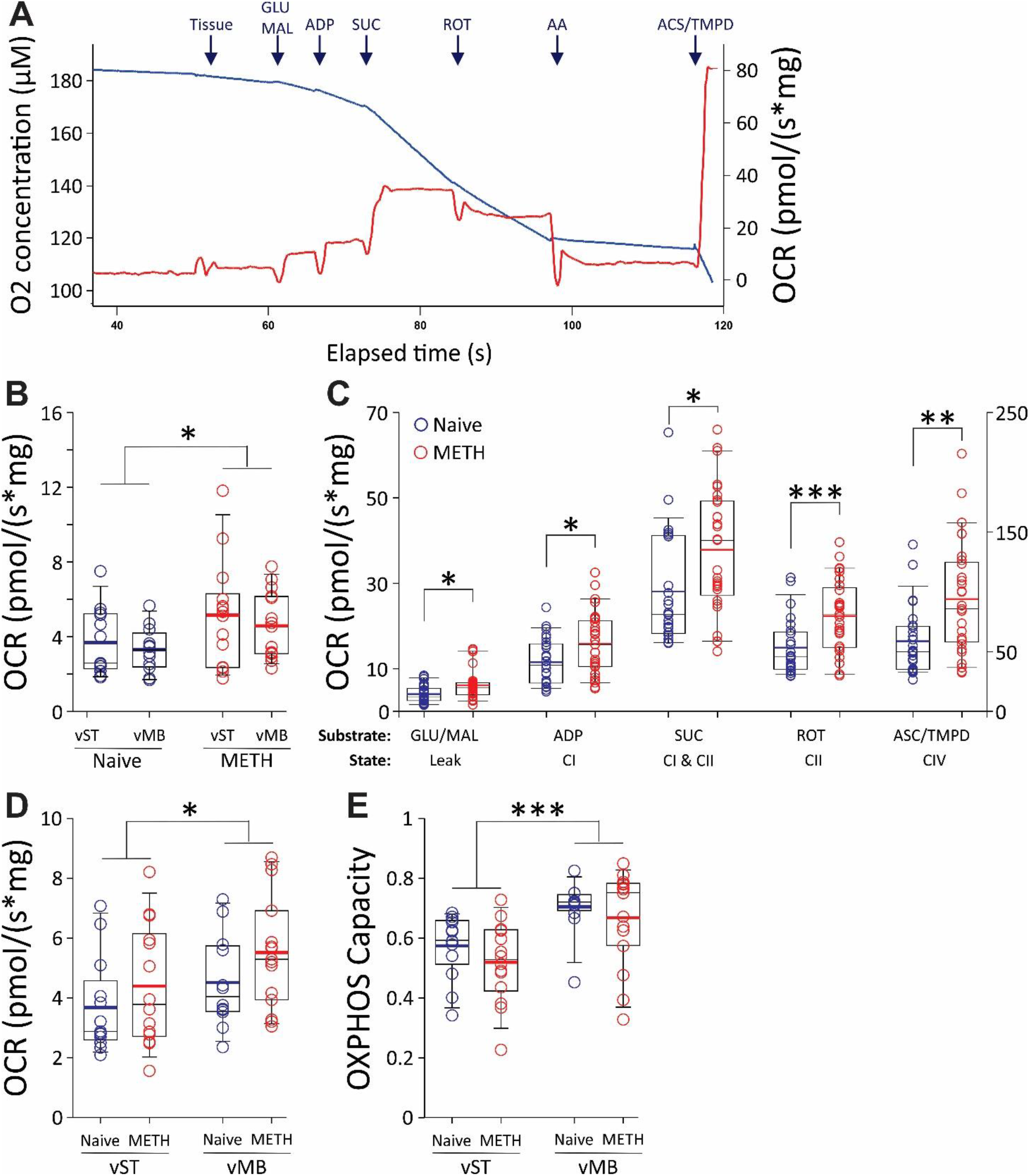
Mitochondrial respiration after methamphetamine (METH) self-administration in mice. A) An example of the protocol used in our experiments shows the oxygen consumption rate (OCR, red line) and the (O2) concentration (blue line). Application of the substrates or inhibitors is shown on the top: Glutamate (GLU), Maleate (MAL), Adenosine diphosphate (ADP), Succinate (SUC), Rotenone (ROT), Antimycin A (AA), Ascorbate (ASC) and N, N, N’, N’-tetramethyl-p-phenylenediamine (TMPD). B) Basal tissue respiration without any reagent was increased after METH self-administration in both ventral striatum (vST) and the ventral midbrain (vMB), compared with tissue from naive animals. C) After correction for no mitochondrial respiration (response after AA application), increased OCR was observed in the tissue from METH exposed mice in all conditions, independently of the brain region. D) The response to the inhibitory effect of AA was less pronounced in vMB samples than in vST. E) Calculated oxidative phosphorylation (OXPHOS) capability was higher in vMB compared with vST tissue, independently of METH exposure status. Bonferroni Student-t test after ANOVAS: * p <0.05, ** p <0.01, *** p<0.001.

In addition, our calculations indicate a higher complex I-linked OXPHOS capacity in vMB samples independently of METH exposure (F(1,50) = 16.088, p < 0.001, Figure 3E). This can be related to a higher energy metabolic demand of dopamine cell bodies versus their axonal terminals due to the maintenance of firing activity (Pacelli et al., 2015; Pissadaki and Bolam, 2013). However, we did not detect any change of OXPHOS capacity for complex II (METH exposure: F(1,50) = 0.46, NS; brain region: F(1,50) = 1.47, NS). As expected, H_2_O_2_ production changed in response to the different mitochondrial complex activities. However, we did not detect changes in the H_2_O_2_ production rates between samples of naive and METH self-administration exposed mice or between brain regions (data not shown). These data suggest that METH-induced oxidative stress within mitochondria function involves the generation of other oxidative molecules than H_2_O_2_.

### METH self-administration alters dopamine neuron firing activity

Finally, since mitochondrial OXPHOS capacity and ATP production can affect dopamine firing activity *in vitro* (Pacelli et al., 2015), we asked whether our observation of higher mitochondrial respiration after METH self-administration relates to the firing rate in the VTA of METH exposed animals. To test this, we obtained *in vivo* single-unit extracellular recordings from the VTA of anesthetized mice. Recording of dopamine neurons firing under these conditions allows the detection of gross changes in dopamine neurotransmission, including endogenous changes due to diurnal activity rhythms or those induced by pharmacological treatments (Dominguez-Lopez et al., 2014; Dremencov et al., 2009; Valenti et al., 2021). We recorded dopamine cells in seven of nine mice exposed to METH self-administration and eight naive mice.

In agreement with our previous findings (Dominguez-Lopez et al., 2021), the average dopamine dell firing rate was lower in the VTA of mice exposed to METH-self-administration (Naive: 3.12 ± 0.6 (Hz); METH: 1.1 ± 0.2 (Hz); t(73) = 3.5, p < 0.001; Figure 4A). Paradoxically, the number of dopamine neurons recorded per track increased in mice with METH exposure (Naive: 2.0 ± 0.24 (neurons per track); METH: 3.6 ± 0.7 (neurons per track); t(26) = 2.38, p = 0.025; Figure 4B). Increased dopamine neuron population activity seems to be a trademark effect of amphetamines derived from a combination of loss of inhibitory input and increased excitatory input into the VTA from other brain regions like the NAc and the prefrontal cortex (Lodge and Grace, 2008; Valenti et al., 2021). Although the percentage of cells firing in bursts was similar in both groups (Naive: 56.25%; METH-SA: 55.81%), all other parameters of burst activity decreased in cells recorded from the METH self-administration group (Figure 4C-G). Finally, METH lifetime intake inversely correlates with firing rate (r = -0.39, p < 0.001), the percentage of spike in burst (r = -0.35, p = 0.025), and directly correlates with the number of neurons per track (r = 0.447, p =0.012). The increase in dopamine population activity provides information on the source of increased mitochondrial respiration since more spontaneously active cells require more energy, increasing mitochondrial metabolism. Our electrophysiology data support the hypothesis that VTA dopamine firing reaches a new basal activity level, adapting to the constant exposure to METH by increasing the number of spontaneously active neurons at a lower firing activity level.

**Figure 4.**
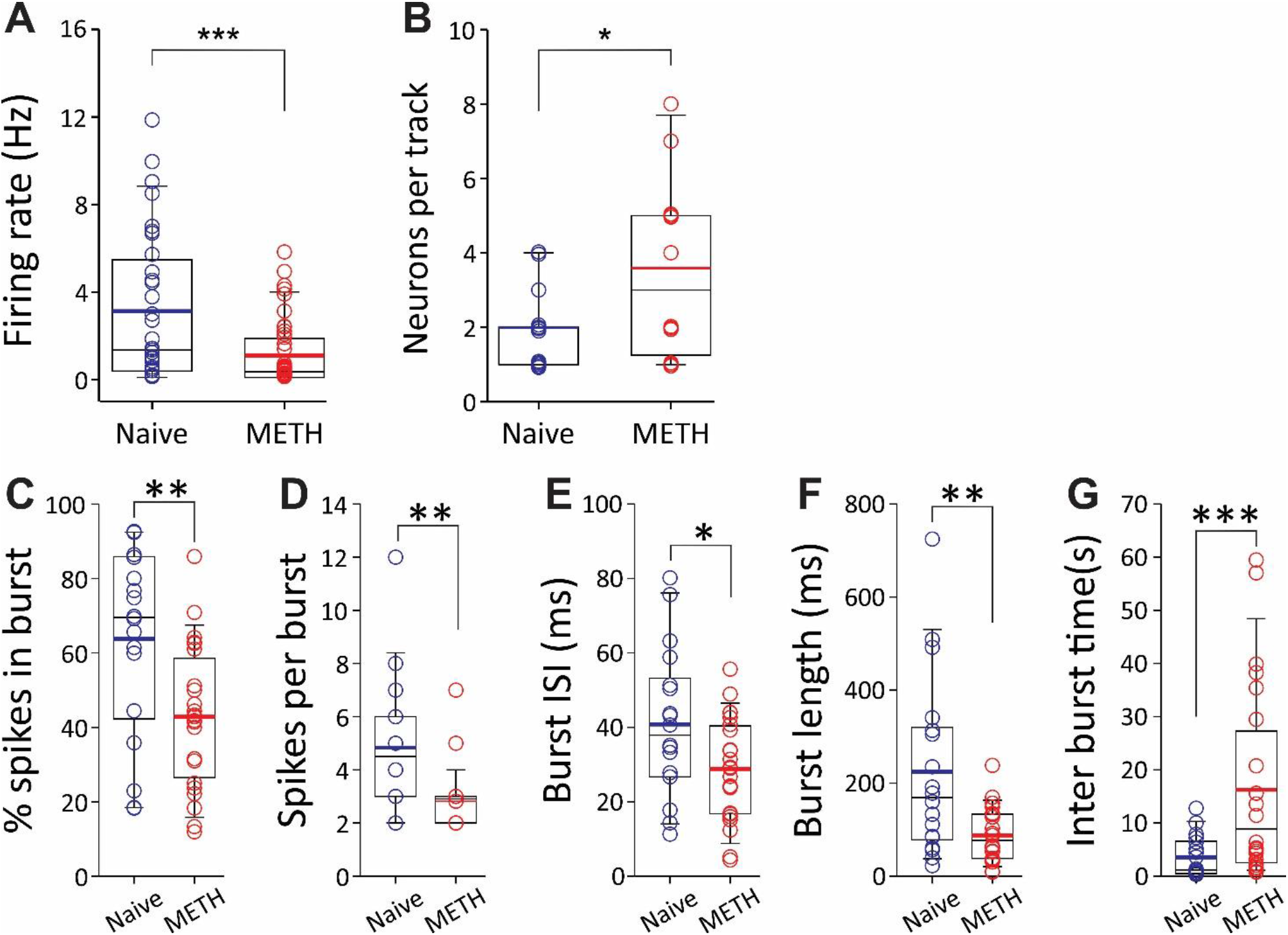
Dopamine neuron firing activity after methamphetamine (METH) self-administration. A) Firing rate from neurons recorded in the ventral tegmental area (VTA) have a lower firing rate in mice exposed to METH self-administration. B) The number of spontaneously active neurons per track was higher in recordings from mice exposed to METH. C-G) Multiple parameters of burst activity in dopamine neurons indicate a decrease in burst activity in mice exposed to METH self-administration. The percentage of spikes fired in a burst, the number of spikes in a burst, the burst interspike interval (ISI), and the burst length decreased, while the time between burst occurrence increased. Simple t-test: * p<0.05, ** p < 0.01, *** p < 0.001.

## Discussion

Our study quantified antioxidant capability and assessed mitochondrial respiration in mesolimbic regions after prolonged exposure to METH self-administration in mice. We also corroborated previous findings on the effect of METH on dopamine neuron firing activity in the VTA. First, our results indicate that the vST, containing dopamine axon terminals, has a higher glutathione antioxidant buffer capability than vMB and declined as METH exposure progressed. Second, mitochondrial respiration increased in vST and vMB after METH self-administration exposure, but only mitochondrial respiration from vMB samples responded to a lifetime intake of METH. Third, surprisingly, we did not detect changes in H_2_O_2_ production due to the increased respiration activity of mitochondrial complexes. Finally, we determined that METH self-administration modified dopamine cell firing rate to reach a new level of basal activity characterized by a higher number of spontaneously active neurons, albeit at lower firing and with decreased burst activity.

### METH self-administration increases striatal oxidative environment

Glutathione is one of the most important antioxidant systems in the brain, responding to oxidative stress by capturing ROS and transforming from GSH (reduced form) to GSSG (oxidized form) (Aoyama et al., 2008). The existing literature suggests that glutathione redox homeostasis is dynamic and responds to METH-induced oxidative stress depending on the exposure time and dose (Achat-Mendes et al., 2007; Harold et al., 2000; Moszczynska et al., 1998). Our study is the first to report that decreased levels of GSH inversely correlate to a lifetime of METH intake in self-administering mice, specifically in vST tissue containing the NAc. Even with relatively low self-administered doses observed in our study, two months of METH self-administration exposure reduced the vST GSH pool, suggesting that this is a progressive pathological condition accompanying METH chronic use (Mirecki et al., 2004). Interestingly, levels of GSH in plasma are high in METH users and remain elevated after two weeks of protracted abstinence (Huang et al., 2013). Since GSH does not cross the blood-brain barrier, the latter observation suggests that outside the brain, the body reacts to METH-induced oxidative stress by increasing GSH levels, but neuronal glutathione synthesis may be limited to compensate for increased oxidative stress (Aoyama et al., 2008).

We also provide the first evidence that in basal conditions, the pool of GSH and its redox capability (GSH/GSSG ratio) are lower in the vMB samples containing the VTA. This observation is likely related to higher axon terminal dopamine release in NAc compared with lower somatodendritic release in the VTA in normal conditions (Iravani et al., 1996). However, after repeated exposure, the ability of METH to induce dopamine release decreases, which could potentially influence its ability to modify GSH pools in the mesolimbic pathway (Zhang et al., 2001). Furthermore, our data indicate that H_2_O_2_ production in vST and vMB does not increase even though mitochondrial respiration does increase following METH self-administration. Therefore, the decline in GSH levels in vST may derive from other sources of oxidative stress, including products derived from dopamine catabolism. In line with this interpretation, systemic administration of other antioxidants such as the non-specific ROS scavenger, N-tert-butyl-alpha-phenylnitrone (PBN) or the superoxide-selective scavenger 4-hydroxy-2,2,6,6-tetramethylpiperidine-1-oxyl (TEMPOL) attenuated METH-induced locomotor activity and acutely decreased METH self-administration in rats (Jang et al., 2017).

### METH self-administration increases mesolimbic mitochondria metabolism

Mitochondria dysfunction is considered another component of the neurotoxic effects of METH because noncontingent administration of METH at high doses disrupts mitochondrial complexes function (Brown et al., 2005; Burrows et al., 2000; Feier et al., 2012). In addition, other studies attribute mitochondrial morphological changes and activation of mitochondria apoptotic function to METH exposure (Cadet and Krasnova, 2009; Yang et al., 2018). Although some data reported a transient increase in ATP consumption and mitochondrial respiratory chain function following acute METH activation of dopamine terminals (Shiba et al., 2011), direct evidence of changes in mitochondrial respiratory function after METH self-administration was not previously reported.

Our study shows that mitochondrial respiration in dopamine mesolimbic regions increases after prolonged self-administration exposure. However, it is difficult to determine if increased mitochondrial respiration in our model reflects mitochondrial dysfunction or a higher metabolism induced by METH. Compared to the neurotoxic studies mentioned above reporting dysfunction of mitochondrial complexes, the daily doses of METH self-administered in our mice are comparatively low (< 2 mg/kg). We can interpret this higher mitochondrial metabolism in the dopamine system as a result of the constant effect of METH increasing dopamine synthesis and release, disrupting vesicular storage and dopamine catabolism. Then, increased mitochondria metabolism in the mesolimbic pathway could also reflect the constant engagement of dopamine circuitry encoding the reinforcing properties of METH (Lodge and Grace, 2008; Valenti et al., 2021; Wang et al., 2021). In this context, the direct correlation of vMB mitochondrial respiration with METH intake can be explained by higher energetic demand due to an increase in dopamine neuron population activity in the VTA, as neuronal circuits are recruited (Lodge and Grace, 2008). We did not detect differences in OCR between vST and vMB samples. However, other experimental manipulations such as the local application of benzodiazepines, differentially affect OCR in NAc and VTA (van der Kooij et al., 2018), which suggests that mitochondrial respiration is enhanced in specific pharmacological activated circuits. In addition, we cannot rule out the contribution of other cell types to mitochondrial respiration in our tissue samples. For example, METH acutely suppressed OCR and ATP production in astrocyte cultures, but after one- or two weeks of exposure ex vivo, METH enhanced both parameters of mitochondria respiration (Borgmann and Ghorpade, 2018).

Our study did not observe an increase in H_2_O_2_ production in mesolimbic tissue samples from mice exposed to METH self-administration. Although a surprising result, it is congruent with a study showing that dopamine metabolism by MAO does not increase cytosolic H_2_O_2_ but leads to increased mitochondrial electron transport activity and ATP production to support phasic dopamine release (Graves et al., 2020). This could be a potential mechanism to explain the increased mitochondrial respiration and increased number of neurons detected in our electrophysiological recordings.

### METH self-administration modifies dopamine neuron firing rate

The firing activity of dopamine neurons in the VTA is considered an indicator of the tone of dopamine neurotransmission (Ford, 2014; Lovinger et al., 2022). We previously reported a decrease in dopamine firing activity in the VTA of mice with similar METH self-administration exposure but with a different strain background (Dominguez-Lopez et al., 2021). Our results are consistent with the inhibitory effect of METH on VTA and SN dopamine neuron firing observed in vivo (Kamata and Kameyama, 1985). In contrast, studies in brain slice and culture preparations report a transient increase in dopamine neuron firing after bath perfusion of low concentrations of METH (Branch and Beckstead, 2012; Lin et al., 2016). We first interpreted our data as an adaptative modification after persistent activation of somatodendritic D2 autoreceptors due to METH-induced extracellular dopamine levels and inhibitory feedback (Ford, 2014; Lovinger et al., 2022; Sulzer, 2011).

Although this partly explains the decrease in firing rate and burst activity, the increase in dopamine neuronal population activity does not entirely fit this explanation. In light of our present study, increased mitochondria respiration could derive from an increased number of spontaneously active neurons, as discussed above. Indeed, mitochondria are a dynamic cell structure that adapts to the energy demands of neuronal activity (Cserep et al., 2018). In addition, our study detected an increased complex I-linked OXPHOS capacity in vMB compared with vST samples. However, this is not modified with METH self-administration, suggesting that VTA dopamine neurons have a limited capacity to increase their energy production in response to METH exposure. A similar conclusion was reached by studying basal mitochondria bioenergetics in SN dopamine neuron cultures (Pacelli et al., 2015).

As mentioned, a recently proposed mechanism described the production of ATP derived from dopamine metabolism by MAO and a subsequent increase in mitochondrial electron transport activity (Graves et al., 2020). Since METH disrupts intracellular monoamine homeostasis by inhibiting the action of MAO (Hotchkiss and Gibb, 1980; Suzuki et al., 1980), an alternative explanation for decreased firing activity would be the loss of this ATP supply. Furthermore, prolonged METH exposure eventually decreases striatal dopamine release (Brennan et al., 2010), which could pressure the dopamine system to reach a new basal activity level, including a higher number of spontaneously active neurons. Recently it has been shown that changes in the tonic activity of dopamine neurons reflect the tracking of reward value over time (Wang et al., 2021). Therefore, the effect of METH self-administration resetting dopamine neurotransmission can be indicative of changes in reward perception with METH prolonged use.

## Conclusion

In this manuscript, we presented evidence that METH self-administration promotes an oxidative environment that depletes GSH levels preferentially at the level of the NAc. We also showed that mesolimbic regions reach a higher mitochondrial respiratory metabolism in response to METH self-administration. However, this higher metabolic state does not increase H_2_O_2_ production at the mitochondrial level. We interpreted the increase in mitochondria metabolism as an adaptative process of mesolimbic circuitry due to the constant pressure of METH exposure to sustain dopamine release. This is also reflected in reduced dopamine cell firing but an increased number of spontaneously active neurons in the VTA. Moreover, OCR is becoming an index of dopamine circuit activation that correlates with behavioral and pharmacological manipulations (van der Kooij et al., 2018). Therefore, our data on increased mitochondria respiration in mesolimbic areas could reflect the engagement of this pathway in mediating the reinforcing and rewarding effects of METH, but also a compensatory mechanism to sustain physiological dopamine function in the brain.

## Acknowledgments

Our work received support from NIH grants to SDL (K99DA049719), to the GeroScience Redox Biology Core, Oklahoma Nathan Shock Center of Excellence in Basic Biology of Aging (5P30AG050911), a Department of Veterans Affairs Merit Award to MJB (IO1-BX005396) and funding from the Presbyterian Health Foundation.

